# Normalization of Mass Spectrometry Data (NOMAD)

**DOI:** 10.1101/105783

**Authors:** Carl Murie, Brian Sandri, Timothy J. Griffin, Christine Wendt, Ola Larsson

## Abstract

**Motivation:** iTRAQ reagent-based mass spectrometry (MS) is a commonly used technology for identification and quantification of proteins in biological samples. Such studies are often performed over multiple MS runs, potentially resulting in introduction of MS run bias that could affect downstream analysis. iTRAQ MS data have therefore commonly been normalized using a reference sample which is included in each MS run. We show, however, that such normalization does not efficiently remove systematic MS run bias. A linear model approach was previously proposed to improve on the reference normalization approach but does not computationally scale to larger data. Here we describe the NOMAD (normalization of mass spectrometry data) R package which implements a computationally efficient ANOVA normalization approach with protein assembly functionality.

**Results:** NOMAD provides the same advantages as the linear regression solution but is more computationally efficient which allows superior scaling to larger sample sizes. Moreover, NOMAD efficiently removes bias which allows for valid across MS run comparisons.

**Availability:** The NOMAD Bioconductor package: www.bioconductor.org

## 1 INTRODUCTION

Proteome wide identification and quantification of proteins is considered pivotal for elucidating mechanisms underlying biological systems and pathological states (Bantscheff *et al.*, 2012; Hassanein *et al.*, 2011). For this purpose tandem mass spectrometry (MS/MS)-based quantification using the iTRAQ isobaric tagging reagent (Ross *et al.*, 2004) was introduced and is currently commonly used (approximately 320 publications in 2014, PubMed) (Wu *et al.*, 2006; Aggarwal *et al.*, 2006). In this technology, proteins are chemically labelled with isobaric chemical tags, producing a sample-specific reporter ion or channel within each MS/MS spectrum, which allows for simultaneous quantitative comparison of protein abundance across multiple samples within a single MS run. Current commercially available isobaric tags are limited to eight samples per run. Therefore larger sample sets need to be divided across several distinct MS runs due to the limited multiplexing power of iTRAQ. As a result, the technology offers challenges during normalization not only due to potential sources of bias from sample preparation but in particular because of the potential bias introduced across multiple MS runs.

At present iTRAQ data produced over multiple MS runs is commonly normalized using a reference approach. Thus, a reference sample is measured in one of the iTRAQ channels to allow for relative quantitation between each of the samples in the query channels to the reference. Such relative measures (commonly referred to as iTRAQ ratios) are assumed to have controlled for bias between MS runs. However, in all data sets we have assessed we have observed strong MS run bias after normalization using the reference approach - which will hamper and potentially disqualify downstream analyses (MS run bias from two data sets generated in two different laboratories are shown in figure 1) (Sandberg *et al.*, 2012; Bhargava *et al.*, 2014). This inability in removing MS run bias could potentially be explained because each sample in an MS run uses the same reference for normalization, MS run bias may persist and even potentially be incorporated during normalization to such a reference. The use of reference samples enforces additional limitations including the subsequent reduction in the number of query samples per MS run and a doubling of the variance due to the calculation of abundance ratios (Kerr *et al.*, 2000).

**Figure 1:**
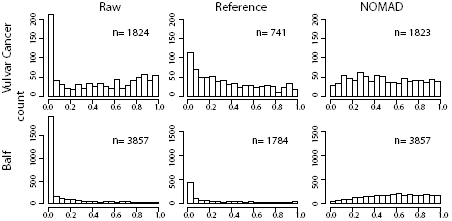
Run Bias. Distribution of p-values produced from testing for a MS run bias on raw, ratio normalized, and NOMAD normalized protein abundance scores. The raw data (first column) show a strong MS run bias which reduces the efficacy of protein abundance comparisons across MS runs. The ratio normalization method reduces the MS run bias but the NOMAD normalization removes the MS run bias entirely.

As a result of these shortcomings, a linear regression approach has been proposed for normalization of iTRAQ data that is applicable to data sets generated over multiple MS runs (Hill *et al.*, 2008; Oberg *et al.*, 2008). Several sources of bias including MS run, iTRAQ label and peptide can be used as regression factors in the model. The residuals of the model are the peptide abundances after removing the bias from the regression factors and these can be used to calculate protein abundances. A significant issue for even relatively modestly sized studies performed across a few MS runs is the computational complexity of solving the linear model resulting in that the method cannot be applied (see results section). Some solutions to this problem have been proposed such as iterative or stagewise regression (Oberg *et al.*, 2008) but to date there is no implementation of this approach. Here we provide the NOMAD (NOrmalization for MAss spectrometry Data) R package implementing an ANOVA normalization method designed for iTRAQ mass spectrometry data in a computationally efficient manner.

## 2 IMPLEMENTATION

The NOMAD package provides two main functions. The first (nomadNormalization) applies an ANOVA model to remove the bias of multiple factors and produces normalized peptide abundances. The second (nomadProteinAsssembly) combines the normalized peptide abundances into summary protein abundances and the user is given multiple options as to how the proteins are assembled. Moreover, the nomadNormalization function allows for correction for imperfect synthesis of isobaric tags. A necessary preprocessing step is to reformat the peptide level output of the MS quantification software (e.g. Protein Pilot) so that it can be used as input in NOMAD. We did not implement a function for such preprocessing (although we provide an example for a commonly used software in the NOMAD R vignette) because of the multitude of tailored software for such quantification.

The structure of the factorial design ANOVA model allows for a simple algebraic solution identical to the more computationally demanding matrix solution of a linear model. We use the following equation for a two-factor ANOVA design including an interaction to illustrate this (Draghici, 2012):

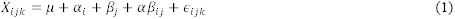

where is *μ* the overall mean and *α* and *β* are factors used in normalization, *i* and *j* are the *i*-th and *j*-th levels of their respective factors, and *k* is the *k*-th data point in the *ij*-th cell. The residuals are defined as:

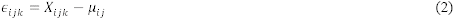

In equation (2) each data point (*X*_*ijk*_) belonging to a particular factor level has the mean of that factor level subtracted (μ_*ij*_). The residual of each data point (∈_*ij*_) is the residual after the means of all factor levels that data point belongs to have been subtracted. This is done in a sequential process where the residuals after subtracting the level means for one factor are used as data for calculation and subtraction of level means of the next factor. This process can be extended to any number of normalization factors and their interactions with the remaining residuals used as the peptide abundances.

The default normalization factors used by NOMAD are the peptide identifiers, protein identifiers, MS run, and iTRAQ channel. We have found that to eliminate the MS run bias the interactions between MS run and the protein, peptide and iTRAQ factors must also be included in the ANOVA model. Users may add or remove any single or interaction factors (in the nomadNormalization function) thereby allowing for custom normalization of in principle any type of data.

Two diagnostic plot functions are included in the NOMAD package. nomadCheckLogTransform generates graphs assessing whether logging is sufficient to produce homogeneous variability and nomadCheckBias produces plots showing the extent of bias from individual factors.

## 3 RESULTS

A key feature of NOMAD is the ability to scale for larger data sets and we therefore compared the performance of the regression approach to NOMAD using data sets of different sizes (table 1). Because NOMAD is both more computationally efficient and scales better than the regression approach, normalization of even modestly sized data sets can only be performed with NOMAD. Moreover, normalization using NOMAD did, in contrast to the reference approach, eliminate MS run bias (figure 1).

**Table 1:** Comparison of computation times between the regression approach and NOMAD. The number (N) of proteins/peptides/data points for each data set tested together with the associated computation times (t) using the linear regression approach (lm function in R) or NOMAD are shown. The data sets (Vulvar) were randomly sampled across 4 MS runs and used 8 iTRAQ channels. Server: 2.00 GHz, 128G RAM, R version: 3.1.2

| N proteins | N peptides | N data points | NOMAD (t) | Lm (t) |
| --- | --- | --- | --- | --- |
| 10 | 435 | 6552 | 0.3s | 44s |
| 100 | 1980 | 32072 | 1.6s | 1hr 31m |
| 200 | 3360 | 53600 | 3s | 9hr 32m |
| 300 | 5046 | 93920 | 7.5s | 63hr 58m |
| 3857 | 72599 | 1144536 | 2m 32s | cannot complete |

## 4 CONCLUSION

NOMAD implements a computationally efficient ANOVA normalization for iTRAQ MS data that scales well for even the largest studies. Moreover NOMAD is better able to address bias from multiple MS runs than the commonly used reference approach. Conveniently, reference samples are not required for NOMAD thus freeing all iTRAQ channels to be used for samples of biological interest. NOMAD thus allows for direct across-MS run comparisons of protein abundances. Therefore, experimental designs can now include more factors of biological interest and increased sample sizes while being normalized efficiently.

## ACKNOWLEDGEMENTS

We would like to thank Janne Lehtiö (Karolinska Institutet) and Ann-Sofi Sandberg (Karolinska Institutet) for sharing raw data. We would like to acknowledge Sue Van Riper (University of Minnesota Center for Mass Spectrpmetry and Proteomics, CMSP) for critical review of this manuscript and Pratik Jagtap (University of Minnesota, CMSP) for technical advice. This research was supported by grants from the Swedish Research Council and the Wallenberg Academy Fellows program (O.L); and NIH R01 HL107612 (C.W.). T.J.G was supported in part by grant 1147079 from the U.S. National Science Foundation.

## REFERENCES

Aggarwal, K. et al. (2006) Shotgun proteomics using the iTRAQ isobaric tags. Brief. Funct. Genomic. Proteomic., 5, 112–120.

Bantscheff, M. et al. (2012) Quantitative mass spectrometry in proteomics: critical review update from 2007 to the present. Anal. Bioanal. Chem., 404, 939–965.

Bhargava, M. et al. (2014) Proteomic profiles in acute respiratory distress syndrome differentiates survivors from non-survivors. PloS One, 9, e109713.

Draghici, S. (2012) Analysis of Variance - ANOVA. Statistics and Data Analysis for microarrays using R and Bioconductor 2nd ed. Chapman and Hill/CRC, Florida, USA.

Hassanein, M. et al. (2011) Advances in proteomic strategies toward the early detection of lung cancer. Proc. Am. Thorac. Soc., 8, 183–188.

Hill, E.G. et al. (2008) A Statistical Model for iTRAQ Data Analysis. J. Proteome Res., 7, 3091–3101.

Kerr, M.K. et al. (2000) Analysis of Variance for Gene Expression Microarray Data. J. Comput. Biol., 7, 819–837.

Oberg, A.L. et al. (2008) Statistical Analysis of Relative Labeled Mass Spectrometry Data from Complex Samples Using ANOVA. J. Proteome Res., 7, 225–233.

Ross, P.L. et al. (2004) Multiplexed Protein Quantitation in Saccharomyces cerevisiae Using Amine-reactive Isobaric Tagging Reagents. Mol. Cell. Proteomics, 3, 1154–1169.

Sandberg, A. et al. (2012) Tumor proteomics by multivariate analysis on individual pathway data for characterization of vulvar cancer phenotypes. Mol. Cell. Proteomics MCP, 11, M112.016998.

Wu, W.W. et al. (2006) Comparative study of three proteomic quantitative methods, DIGE, cICAT, and iTRAQ, using 2D gel- or LC-MALDI TOF/TOF. J. Proteome Res., 5, 651–658.

